# Endosomal cAMP production broadly impacts the cellular phosphoproteome

**DOI:** 10.1101/2021.01.06.425636

**Authors:** Nikoleta G. Tsvetanova, Michelle Trester-Zedlitz, Billy W. Newton, Grace E. Peng, Jeffrey R. Johnson, David Jimenez-Morales, Andrew P. Kurland, Nevan J. Krogan, Mark von Zastrow

## Abstract

Endosomal signaling downstream of G protein-coupled receptors (GPCRs) has emerged as a novel paradigm with important pharmacological and physiological implications. Yet, our knowledge of the functional consequences of intracellular signaling is incomplete. To begin to address this gap, we combined an optogenetic approach for site-specific generation of the prototypical second messenger generated by active GPCRs, cyclic AMP (cAMP), with unbiased mass spectrometry-based analysis of phosphoproteomic effects. We identified 218 unique, highconfidence sites whose phosphorylation is either increased or decreased in response to cAMP production. We next determined that the same amount of cAMP produced from endosomes led to more robust changes in phosphorylation than the plasma membrane. Remarkably, this was true for the entire repertoire of identified targets, and irrespective of their annotated sub-cellular localization. Furthermore, we identified a particularly strong endosome bias for a subset of proteins that are dephosphorylated in response to cAMP. Through bioinformatics analysis, we established these targets as putative substrates for protein phosphatase 2A (PP2A), and we propose compartmentalized activation of PP2A-B56δ as the likely underlying mechanism. Altogether, our study extends the concept that endosomal signaling is a significant functional contributor to cellular responsiveness by establishing a unique role for localized cAMP production in defining categorically distinct phosphoresponses.

## Introduction

Cells dynamically respond to their surrounding through the precise actions of transmembrane receptors, among which G protein-coupled receptors (GPCRs) comprise the largest and most versatile class. GPCRs are seven transmembrane receptors which control all essential mammalian physiology, and have become key targets for pharmacological intervention for human diseases^1^. In a classical GPCR cascade, a ligand-bound receptor stimulates adenylyl cyclase via Gαs to produce the second messenger cyclic AMP (cAMP). cAMP directly modulates a handful of effectors, including the prototypical protein kinase A (PKA), to promote a plethora of changes in the cellular environment. Receptors were long presumed to transduce their entire repertoire of signaling consequences via Gαs/cAMP from the cell surface. Recently, however, it has become clear that GPCRs can be activated to generate cAMP from endosomal membranes as well^2^, and accumulating evidence suggests that this compartmentalization can underlie unique physiology and selective drug responses^3,4^. However, our understanding of the mechanisms that give rise to these spatially biased outcomes remains very limited.

Here, we apply a reductionist approach to ask whether production of the same second messenger, cAMP, from distinct sites can yield discrete outcomes. Since the initial signal is propagated downstream through the dynamic interplay between protein-protein interactions and post-translational modifications, we reason that dissection of the spatial regulation of these early steps could yield important insights into how compartmentalized signaling may operate to uniquely rewire the cell. To do so, we carry out a comprehensive survey of the phosphoproteomic changes driven by localized generation of cAMP. We circumvent the technical challenges associated with manipulating the localization of transmembrane receptors by utilizing a previously validated optogenetic strategy^5^to generate the relevant proximal signal, cAMP, from the plasma membrane or early endosomes under matched photostimulation conditions. By coupling this approach with unbiased analysis using quantitative mass spectrometry, we show that endosomal cAMP exerts location bias on a wide range of downstream phosphoresponses.

## Results and Discussion

We recently described an optogenetic approach to enable spatiotemporal control over cAMP generation, which utilizes a bacteria-derived photoactivatable adenylyl cyclase, bPAC, fused to organelle-specific targeting sequences^5,6^. By transiently expressing bPACs localized either on early endosomes (“bPAC-Endo”) or on the plasma membrane (“bPAC-PM”), we reported that endosomal but not plasma membrane cAMP production gave rise to robust downstream transcriptional responses^5^. Here, we set off to apply this strategy to ask whether there is similar selectivity based on location for the cellular cAMP-dependent phosphoresponses.

In order to reduce variability that could arise from non-uniform bPAC expression in transiently transfected cells, we first generated clonal HEK293 cell lines stably expressing the following optocyclase constructs: 1) “bPAC-PM”, 2) “bPAC-Endo”, and 3) cytosolic bPAC (“bPAC-Cyto”) to serve as non-localized control **(Fig. S1a)**. We verified expression and appropriate subcellular localization of the constructs by flow cytometry **(Fig. S1b)**, and fluorescence microscopy **(Fig. 1a)**, respectively. In addition, we confirmed that photoactivation of bPAC-Endo and bPAC-Cyto in the stable cell lines drove more robust transcriptional upregulation of an established cAMP target gene than photoactivation of bPAC-PM **(Fig. S1c)**, as previously observed in cells transiently expressing bPACs^5^. Next, we used stable isotope labeling with amino acids in cell culture (SILAC) and mass spectrometry in order to globally identify and quantify proteins phosphorylated in response to cAMP production **(Fig. S1a)**. bPACs were activated with light for 5 min, cells incubated for an additional 5 min in a dark incubator, and then subjected to lysis. Next, proteins were extracted and digested, phosphorylated peptides were isolated by iron(III)-nitrilotriacetic acid immobilized metal ion affinity chromatography and subjected to liquid chromatography-tandem mass spectrometry. Unstimulated cells served as negative control, and SILAC media were swapped for each cell line to rule out non-specific effects. Media swapped samples were considered biological replicates. The maximum intensity of any unique peptide and charge state between the two technical replicates were used to condense the data and yield one value per peptide per biological replicate.

**Figure 1.**
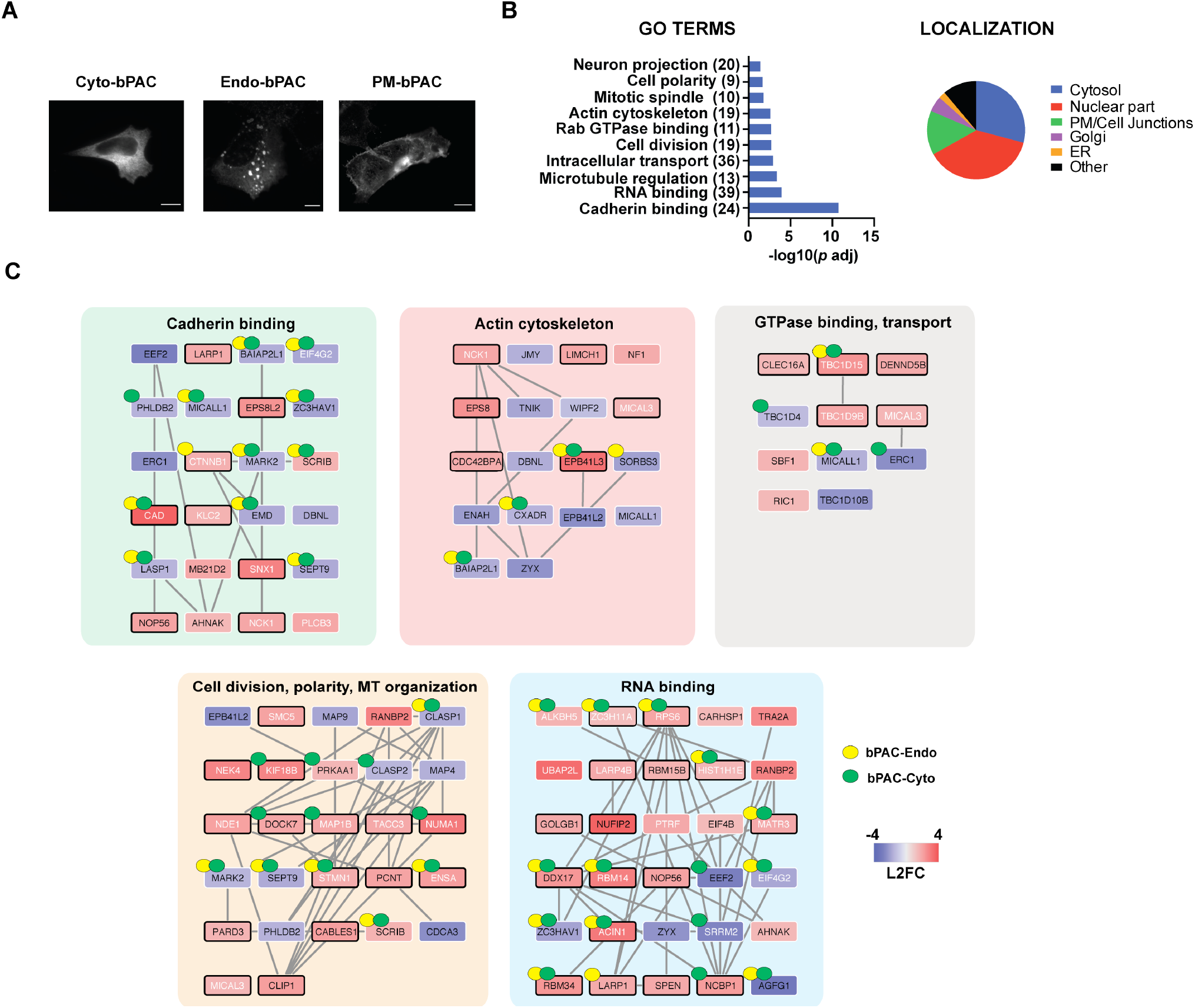
cAMP signaling induces profound changes in the cellular phosphoproteome. **(a)** Subcellular localization of bPAC constructs visualized by immunofluorescence microscopy with Alexa-conjugated anti-myc antibody in fixed cells. PM = plasma membrane, Endo = early endosome, Cyto = cytosol. Scale bar = 10 μm. **(b)** Gene Ontology (GO) categories and annotated subcellular localization of proteins with significant changes in phosphorylation upon cAMP production **(Dataset S1)**. The number of unique target phosphoproteins identified in the mass spectrometry analysis that belong to each category is shown in parentheses. **(c)** Proteins in each GO category were mapped to known interaction networks using StringDB and visualized in Cytoscape (see “Materials and Methods” for additional details). The interactors are color-coded based on their phenotype as indicated in the legend (red = increased phosphorylation, blue = decreased phosphorylation, shade = magnitude of change). White node font label indicates sites previously shown to change in phosphorylation status upon cAMP induction, and black font designates novel sites identified in this study. Black border color indicates the presence of a predicted or known PKA-binding motif. Yellow and green dots = sites significantly biased towards regulation from endosomal or cytosolic cAMP relative to plasma membrane cAMP **(Dataset S2)**.

We first focused on sites and proteins phosphorylated in response to photostimulation in the bPAC-Cyto samples, as these would presumably reflect targets of bulk cAMP and be independent of the subcellular site of signal origin. Across the biological replicates, we obtained quantitative information for 9,285 phosphopeptides corresponding to 2,227 unique proteins. Of these, 6,035 peptides (1,946 proteins) were quantified in both replicates with significant degree of reproducibility (Pearson coefficient = 0.76, *p* < 1.0×10^−4^) **(Fig. S2a, Dataset S1)**. To identify highconfidence targets, we considered only phosphopeptides that displayed abundance changes in response to photostimulation with z-scores ≥ 2 or ≤ −2 in each replicate. Under these cut-off criteria, 218 unique sites within 232 phosphopeptides derived from 184 proteins were classified as cAMP target sites (Pearson coefficient = 0.99, *p* < 1.0×10^−4^). Of these 232 phosphopeptides, 168 peptides increased and 64 decreased in abundance relative to matched untreated controls reflecting increase or decrease in phosphorylation status, respectively **(Dataset S1)**. Comparison with available phosphoproteomic datasets derived from studies of GPCR/cAMP activation revealed extensive overlap (65/218 sites = 30%)^7–14^. Of particular significance, we confirmed 1/3 of GPCR/cAMP target phosphosites previously reported in the same cell type, HEK293 cells^14^. Further consistent with signaling through the canonical cAMP-PKA axis, a large fraction of the modified peptides (107/232 = 46%, *p* < 1.0×10^−6^ by Fisher’s exact test) contained the basophilic sequence motif R-R/K-X-p(S/T), which corresponds to the known PKA target site^15^. Taken together, these data strongly suggest that the remaining novel sites likely represent bona fide phosphotargets of the cascade.

To gain insight into the biological pathways and downstream processes impacted by cytosolic cAMP, we performed Gene Ontology (GO) and network analyses on the modified proteins. These analyses revealed a prevalence for a number of pathways known to be associated with the cAMP cascade, including cell adhesion, alteration of actin cytoskeletal dynamics, changes in cellular architecture via modulation of GTPase activity, and interplay with intracellular cargo trafficking^16–18^ **(Fig. 1b, Dataset S1)**, providing further validation for the phosphoproteomic dataset. The proteins within these categories localize to different sub-cellular compartments (nucleus, cytosol, focal adhesions, plasma membrane, and intracellular vesicles; **Dataset S1**), and network analysis found that a large fraction of these proteins participate in shared complexes **(Fig. 1c)**. In addition to the known effects of cAMP production, we also observed changes in the phosphorylation status for a significant number of RNA processing proteins, a GO category that to our knowledge has not been seen in proteomics studies of this pathway before. While cAMP is known to exert pronounced effects on cellular gene expression via PKA-dependent modulation of transcription factors, such as the cAMP response element-binding protein CREB, the impact of this cascade on post-transcriptional regulation of gene expression is vastly underexplored. Yet, RNA-binding proteins constituted the largest fraction of hits identified here (39/184 = 21%, **Fig. 1b-c**, **Dataset S1**), highlighting an avenue of potential significance for future exploration.

Using the defined set of cAMP target peptides derived from the bPAC-Cyto experiments, we next asked if the subcellular site of second messenger origin impacts these responses. We first carried out unbiased hierarchical clustering analysis across all experimental conditions, which revealed higher degree of similarity between the signaling profiles of bPAC-Cyto and bPAC-Endo cells, and a different signature in bPAC-PM cells **(Fig. 2a)**. To dissect the nature of these differences, we next performed pairwise comparisons of cAMP target peptide abundance between experimental conditions. This analysis showed that cAMP originating at the plasma membrane gave rise to less robust changes in phosphorylation compared to the cytosol or the endosome. The trend held for all protein targets, regardless of their respective sub-cellular localization or whether they were decreased or increased in phosphorylation upon cAMP generation **(Fig. 2b, Fig. S2b)**. Further, we found that the apparent bias towards endosomal over plasma membrane-derived cAMP was statistically significant in the case of 64 peptides corresponding to 62 unique proteins; and similarly, 86 peptides derived from 81 proteins were selective for cytosolic cAMP (FDR = 0.1 by multiple t-test analysis, **Fig. 2c, S2c**, **Dataset S2**). In contrast, we detected no statistically significant differences in the phospho-signatures between the bPAC-Cyto and bPAC-Endo samples **(Fig. S2c, Dataset S2)**. To corroborate that these results were not biased by inputting a set of pre-defined cAMP targets based on the bPAC-Cyto experiments, we applied the same hit-calling criteria (see above) to identify phosphotargets in either bPAC-Endo or bPAC-PM stimulated conditions, respectively, and repeated the pairwise analyses. The same trends held across these comparisons (compare **Fig. 2b and Fig. S2d)**. Thus, phosphoresponses to cAMP originating from endosomes and the bulk cytosol closely resemble each other and are more robust than phosphoresponses triggered by cAMP coming from the cell surface.

**Figure 2.**
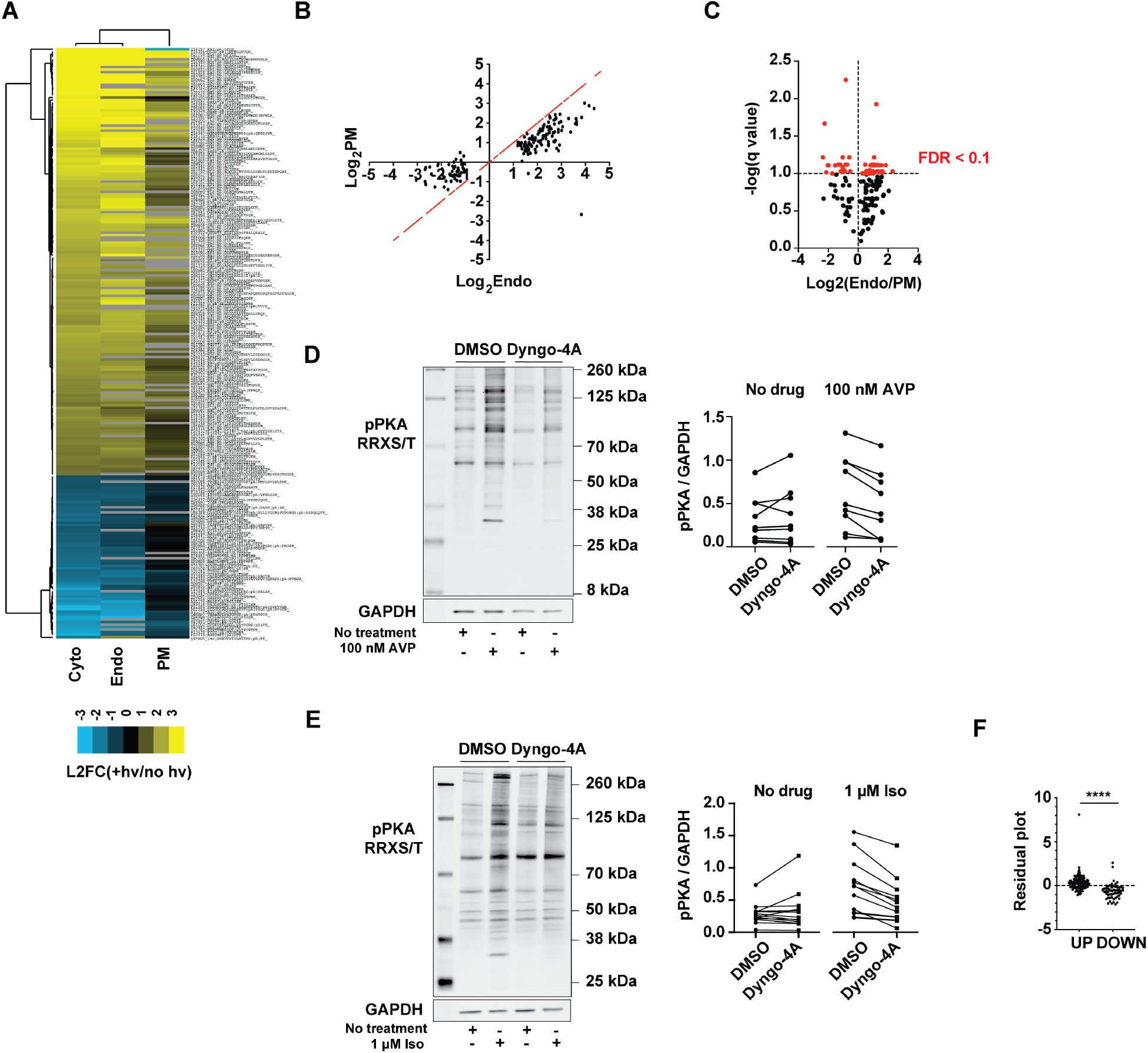
Spatial bias of the cAMP-dependent phosphoproteomic responses. **(a)** Heat map showing the changes in abundance of cAMP-dependent phosphotargets (indicated on the y-axis) averaged across biological replicates for each of three experimental conditions (x-axis), relative to the uninduced controls. Peptides and conditions were subjected to average linkage hierarchical clustering analysis using Euclidian distance as a similarity metric. Yellow = upregulated, blue = downregulated, black = unchanged, grey = peptides with missing values. **(b)** Scatter plot comparing the abundance of cAMP target phosphopeptides in bPAC-Endo and bPAC-PM cells. Red dotted line denotes x = y. **(c)** Volcano plot of differentially modified peptides in bPAC-Endo versus bPAC-PM cells. Significant peptides with FDR ≤ 10% are color-coded in red and summarized in **Dataset S2**. Data in **(a-c)** are average of n = 2. **(d-e)** Western blot analysis of phospho-PKA substrate abundance (pPKA RRXS/T) in cells upon activation of V2R **(d)** or β2-AR **(e)** signaling with or without GPCR endocytic blockade. Cells were pre-treated with 30 μM Dyngo-4a or vehicle (DMSO) for 15 min, then stimulated with 100 nM AVP **(d)** or 1 μM isoproterenol **(e)** for 10 min. Representative anti-phospho PKA blots are shown. Quantification of individual phospho-PKA substrate bands relative to GAPDH (for bands included in the analyses, see **Fig. S3b,d**) are average from n = 3 experiments. **(f)** Plot of residuals computed from linear regression analysis on bPAC-Endo and bPAC-PM mass spectrometry datasets. Residuals are shown for upregulated and downregulated phosphopeptides. Data are average of n = 2. **** = *p* ≤ 0.0001 by two-sided unpaired t-test analysis. PM = plasma membrane, Endo = early endosome, Cyto = cytosol.

Because bPACs are an engineered system to generate cAMP, we aimed to provide initial validation for these observations using relevant GPCR models known to stimulate cAMP production from the plasma membrane and early endosomes. We first focused on the vasopressin receptor, V2R, which generates sustained cAMP responses from endosomal membranes **(Fig. S3a)** to regulate water homeostasis^19^. We measured endogenous PKA phospho-substrate abundance following cAMP stimulation through transiently expressed V2Rs by Western blot analysis, and detected robust increase in PKA-dependent phosphorylation in cells treated with vasopressin **(Fig. 2d, S3b)**. In order to parse out the contribution of each signaling compartment to PKA activity, V2R endocytosis was blocked by acute inhibition of dynamin with the drug Dyngo-4A^20^. When V2R activation was confined to the plasma membrane through this manipulation, we saw blunted cAMP production and phosphorylation of PKA substrates after addition of vasopressin **(Fig. 2d, S3a-b)**. Next, we examined localized phosphosignaling from the beta2-adrenergic receptor (β2-AR), which is expressed endogenously in our cell line. In contrast to the V2R, cAMP signaling from the β2-AR is transient, and the effect of endosomal activation on global cAMP production is small **(Fig. S3c)**^5,21,22^. Yet, intracellular β2-ARs were shown to be essential for eliciting downstream transcriptional responses^5,22,23^. Similarly to what we observed with overexpressed V2R, β2-AR stimulation led to increase in PKA substrate phosphorylation, which required receptor internalization **(Fig. 2e, S3d)**. Thus, these data provide additional support to the bPAC phosphoproteomic results by showing a requirement for GPCR internalization and endosomal signaling for successful activation of PKA and regulation of its downstream substrates for two receptors with distinct endosomal cAMP signaling responses.

Among the spatially biased proteins, we found a number of important intracellular effectors including several essential kinases **(Table 1)**. Because these enzymes can in turn give rise to an array of downstream cellular responses, we reasoned that one of these location-biased master regulators could underlie the divergent signaling profiles observed in bPAC-Endo and bPAC-PM cells. Further inspection of the trends in abundance between the downregulated and upregulated target sites gave us a hint regarding the identity of this regulator. We saw that the set that increased in phosphorylation in response to cAMP was modified under all conditions, albeit more robustly by endosomal than plasma membrane cAMP **(Fig. 2b-c)**. In contrast, we noticed that a large number of sites were dephosphorylated *only* by bPAC-Endo (**Fig. 2b-c)**. This suggested that the downregulated phosphosites might be disproportionally impacted by the location of cAMP origin. To test this possibility, we performed linear regression analysis comparing the abundance values between the bPAC-PM and bPAC-Endo samples, computed the residuals for the regression, and used unpaired t-test to determine if the distributions of residual values were significantly different between the upregulated and downregulated sets of phosphosites. This analysis indicated that the abundance of dephosphorylated peptides was indeed more significantly biased by location (*p* < 1.0×10^−4^ by unpaired t-test) **(Fig. 2f, Fig. S3e)**.

**Table 1.** Kinases differentially regulated by cAMP localization.

| Uniprot ID | Gene | Description | Modified Residue | Endo (± SD) | PM (±SD) |
| --- | --- | --- | --- | --- | --- |
| P04049 | Raf1 | RAF proto-oncogene Ser/Thr-protein kinase; a regulatory link between the Ras GTPases and the MAPK/ERK cascade in determining cell fate decisions including proliferation, differentiation, apoptosis, survival and oncogenic transformation. | Ser43 | $2.11 \pm 0.06$ | $1.60 \pm 0.11$ |
| Q00536 | Cdk16 | Cyclin-dependent kinase 16; plays a role in vesicle-mediated transport processes and exocytosis. | Ser153 | $2.51 \pm 0.03$ | $1.79 \pm 0.15$ |
| Q9BZ23 | Pank2 | Pantothenate kinase 2, master regulator of the CoA biosynthesis. | Ser189 | $1.51 \pm 0.15$ | $0.74 \pm 0.01$ |
| Q7KZI7 | Mark2 | Ser/Thr-protein kinase MARK2; regulates cell polarity and microtubule dynamics. | Ser619 | $-1.35 \pm 0.04$ | $-0.47 \pm 0.19$ |

To dissect the mechanistic basis for these differences, we next asked how these sites are regulated. While our initial motif analysis done on the entire set of 232 cAMP target peptides revealed a single enriched motif, the PKA preferred site, R-R/K-X-(S/T), we reasoned that this analysis might have been skewed disproportionally by the prevalence of upregulated relative to downregulated peptides (168 vs 64 peptides, respectively). Therefore, we went back and searched for motifs enriched only among the 64 phosphopeptides decreased in abundance, and found that this group was dominated by X-p(S/T)-P, where P is proline (*p* < 1.0×10^−6^ by Fisher’s exact test). While ~70% of all depleted phosphotargets had this motif, the incidence of X-p(S/T)-P was even more striking when we narrowed down the list to include only the peptides preferentially dephosphorylated in response to endosomal relative to plasma membrane cAMP **(Dataset S2)**. In this subset, 20 of these 22 peptides (>90%) had the motif (*p* < 1.2×10^−4^ by Fisher’s exact test). Two protein phosphatases, PP1A and PP2A, are known to modify proline-directed sequences^11,24^. Hence, we searched the mass spectrometry data and found a peptide corresponding to a known regulatory site within the PP2A-B56δ subunit, S573. PP2A exists as a heterotrimer, composed of a dimeric core, the catalytic (C) and structural (A) subunits, and a regulatory (B) subunit, which controls the localization and substrate specificity of the enzyme. Serine 573 within B56δ is an established target of PKA-dependent phosphorylation that is required for PP2A enzymatic activity^25,26^. While the S573-containing phosphopeptide had not passed the stringent cut-off criteria used to define the set of cAMP targets, it was reproducibly and robustly upregulated (>1.5-fold) by both cytosolic and endosomal cAMP, but not by second messenger generated at the plasma membrane **(Fig. 3a)**. This location bias in pS573 abundance is consistent with the trends seen in protein dephosphorylation between experimental conditions **(Fig. 2a-b, Fig. S2b)**, and therefore supports differential PP2A activity as the likely underlying mechanism.

**Figure 3.**
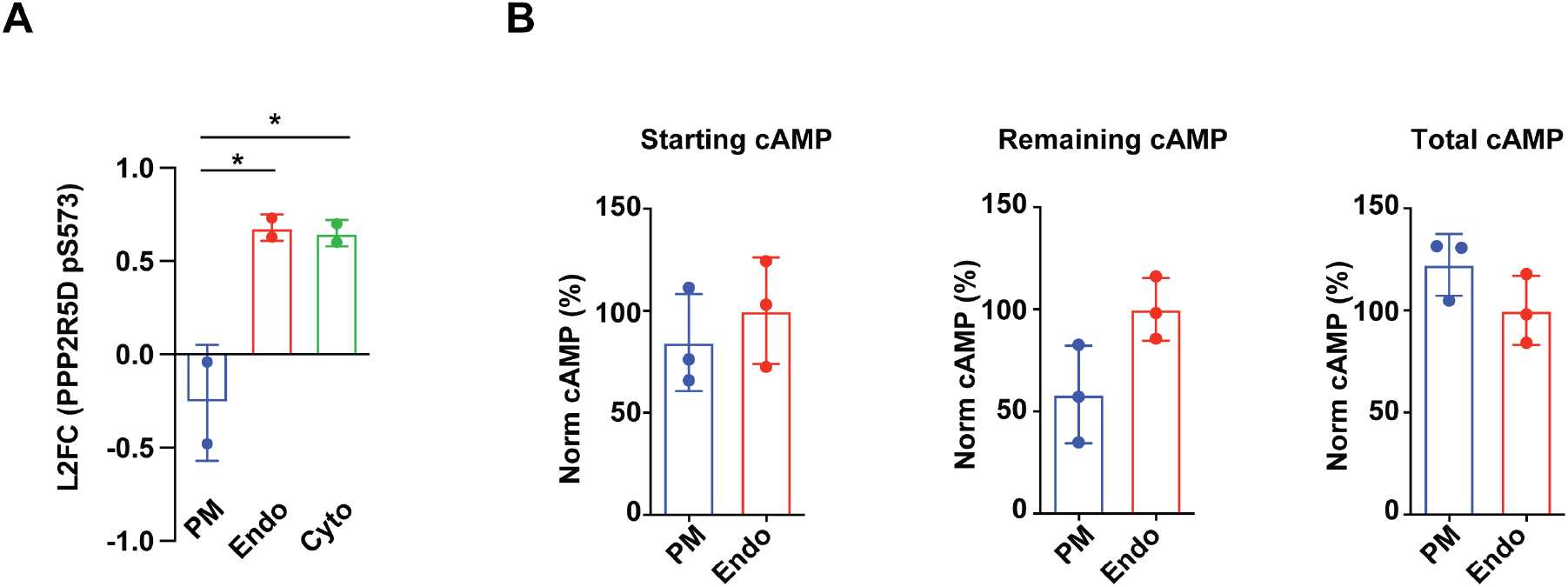
Location-specific phosphatase activation and cAMP hydrolysis as mechanisms underlying spatial encoding of the cAMP-dependent phosphoresponses. **(a)** cAMP signaling promotes changes in the abundance of a known regulatory site in PP2A-B56δ. Fold changes in phosphopeptides abundance were quantified by SILAC-mass spectrometry analysis. Data are mean of n = 2. Error bars = ± SD. **(b)** cAMP accumulation was measured by ELISA assay directly after light stimulation (“Starting cAMP”, left panel) or after light stimulation and dark incubation in the absence (“Remaining cAMP”, middle panel) or presence of the phosphodiesterase inhibitors, 100 μM IBMX and 10 μM Rolipram (“Total cAMP”, right panel). All cAMP measurements were normalized to the total protein concentration of each respective sample, and are displayed as percentage of the mean cAMP amount from the bPAC-Endo samples. Data are mean of n = 3. Error bars = ± SD. * = *p* ≤ 0.05 by one-way ANOVA test, using Tukey’s multiple testing correction.

That cAMP originating at the plasma membrane gives rise to less robust changes in transcription^5^ and phosphorylation underscores the existence of a “barrier” preventing cAMP produced at the cell periphery from transducing its full repertoire of downstream effects. One known mechanism for establishing spatial gradients of second messenger is through local hydrolysis by phosphodiesterase enzymes (PDEs)^27^, which prompted us to examine if there are differences in the kinetics of cAMP degradation between the plasma membrane and the endosome. To this end, we measured cAMP concentrations under three conditions matching the timecourse of the mass spectrometry experiment: 1) immediately following a 5-min photostimulation, which reflects peak induced amounts (“starting cAMP”), 2) after 5 min of photostimulation followed by 5 min in a dark incubator (“remaining cAMP” after degradation), and 3) after 5 min of photostimulation followed by 5 min in a dark incubator when cells pre-treated with the PDE inhibitors rolipram and IBMX to prevent hydrolysis (“total cAMP generated”). Quantification of starting cAMP showed that, at matched doses of light stimulation, bPAC-PM and bPAC-Endo produce comparable amounts of the second messenger **(Fig. 3b, left panel)**. However, by the end of the 10 min timecourse, the “remaining cAMP” in the bPAC-PM cells was half the amount measured in bPAC-Endo cells **(Fig. 3b, middle panel)**, suggesting that bPAC sub-cellular localization impacts the duration of the resulting cAMP signal. As this phenotype was reversed by PDE inhibition **(Fig. 3b, right panel)**, the results are consistent with higher cAMP hydrolysis in the vicinity of the plasma membrane. Therefore, we propose that differential cAMP hydrolysis between compartments could be a contributing factor to reduced efficiency of phosphosignaling to cytoplasmic and nuclear substrates.

The present work establishes a unique role for cAMP signaling location in defining categorically distinct cellular phosphoresponses, and it provides a mechanistic framework for this functional specialization paradigm. Through global phosphoproteomics analysis, we generate an extensive cellular atlas of cAMP-modulated proteins that, collectively, participate in all essential aspects of cellular physiology **(Fig. 1)**. We show that the location of cAMP origin impacts the regulation of the *entire repertoire* of targets regardless of sub-cellular localization, including a number of important intracellular signaling effectors **(Fig. 2, Table 1)**. We further interrogate the responses downstream of two Gαs-coupled GPCRs, the V2R and β2-AR, known to generate cAMP from the plasma membrane and endosomes^5,19,21^. We find that phosphosignaling through PKA, a canonical cAMP-dependent kinase, is blunted under endocytic blockade, concurrent with diminished cAMP production **(Fig. 2, S3)**. These results mirror the general trends toward more efficient activation of these cascades from the endosome observed with the optogenetic system. We note, however, that under the experimental conditions used in this study, total cAMP production from localized bPACs was more robust than following agonist stimulation of the V2R **(Fig. S3f)** or β2-AR^5^. Therefore, the degree of overlap between the phosphotargets and pathways regulated downstream of the optogenetic system or via receptors remains to be determined. Through bioinformatics analysis, we further find that proteins containing a PP2A consensus motif are disproportionately biased by the site of signaling, which reflects an apparent inability of the plasma membrane-derived signal to drive PKA-dependent phosphorylation and activation of PP2A **(Fig. 3)**. PP2A heterotrimers containing the B56δ subunit are highly enriched in the nucleus^28,29^, and not surprisingly the majority of proteins dephosphorylated in a location-dependent manner concentrate predominantly or partially in the nucleus by annotation **(Datasets S2)**. This observation is even more striking in light of our recent discovery that another nuclear cAMP effector and cellular master regulator, CREB, is similarly activated by PKA only in response to cytosolic and endosomal signals^5^. Taken together, these findings reinforce a unique functional coupling between endosomal cAMP and nuclear target activation that warrants further investigation. We propose that the unique ability of endosomal cAMP to robustly stimulate downstream responsiveness arises in part from higher cAMP degradation rates at the plasma membrane **(Fig. 3)**. Whether this mechanism alone is sufficient to fully account for the location bias of phosphosignaling or if there also exist endosome-specific effectors enhancing the selectivity remains to be determined in future studies.

While the optogenetic strategy outlined here enabled us to directly assess the downstream responses to organelle-based cAMP production, the approach is engineered and may not fully capture the breadth of signaling outcomes generated by localized GPCRs. cAMP responses from compartmentalized bPACs were induced under a single matched stimulation condition from two known signaling locations to demonstrate that, in principle, production of the same second messenger from distinct sites can yield discrete outcomes. Yet, GPCRs vary in the amounts and kinetics of plasma membrane- and endosome-derived cAMP production, and their diverse spatiotemporal dynamics could further fine-tune the responses. Our experimental strategy can be applied to investigate this aspect by varying the duration of photostimulation of localized bPACs. Additionally, GPCRs signal through a multitude of downstream effectors, including G proteins, topologically distinct cAMP sources (transmembrane and soluble adenylyl cyclases)^30^, and arrestins^3^. With an ever-growing list of compartmentalized receptor cascades, distinct second messengers (cAMP, cGMP, calcium), and additional intracellular membranes being ascribed roles in signaling across cell types, we anticipate that there will be further layers of complexity and functional specificity to be explored.

## Experimental Procedures

### Chemicals and antibodies

(-)-Isoproterenol hydrochloride was purchased from Sigma-Aldrich (Cat #I6504), dissolved in water/100 mM ascorbic acid to 10 mM stock, and used at 1 μM final concentration. Arginine vasopressin acetate salt was purchased from Sigma-Aldrich (Cat #V9879), dissolved in water to 1 mM stock, and used at 100 nM final concentration. Dyngo-4a (AbCam, Cat #ab120689) was dissolved in DMSO to 30 mM, stored protected from light and added to cells to 30 μM final concentration in serum-free DMEM. Protease inhibitor cocktails were purchased from Roche (Cat #04693159001, 11836170001) and used according to manufacturer recommendations. Phosphodiesterase inhibitors, IBMX (3-Isobutyl-1-Methylxanthine) purchased from Sigma-Aldrich (Cat #I5879) and Rolipram from Tocris (Cat #0905), were dissolved in ethanol to make 100 mM and 10 mM stocks, respectively. Alexa 647-conjugated mouse anti-myc antibody (Cell Signaling, Cat #2233S) was used at 1:50. Rabbit anti-phospho-PKA substrate (100G7E) antibody was purchased from Cell Signaling (Cat #9624) and used at 1:1000. Mouse anti-GAPDH antibody was purchased from Millipore (Cat #MAB374) and used at 1:1000. Secondary IRDye antibodies were purchased from Li-COR Biosciences and used at 1:10,000 in Odyssey Blocking Buffer (Li-COR Biosciences, Cat #927-50000).

### Cell culture and stable bPAC cell line generation

Human embryonic kidney (HEK293) cells were obtained from ATCC and grown in a CO_2_- and temperature- controlled incubator and propagated in DMEM with high glucose, no sodium pyruvate supplemented with 10% FBS. For V2R experiments, HEK293 cells were transiently transfected with receptor cDNA for 48 h. For SILAC experiments, cells were grown in DMEM deficient in L-arginine and L-lysine and supplemented with L-lysine and L-arginine, or doubly labeled ^13^C-labeled lysine and ^13^C,^15^N-labeled arginine to a final concentration of 0.46 mM each with 10% FBS (Thermo Scientific Cat #89983). Cells were maintained in specific isotope conditions for a minimum of six doublings, with frequent medium changes. To generate clonal cell lines stably expressing bPAC constructs, HEK293 cells were plated on 6-well dishes at 80% confluency, transfected with myc-tagged bPAC constructs under a CMV promoter^5^, plated sparsely the day after transfection to yield single colonies, and selected with 200 μg hygromycin for ~1 month. To quantify bPAC expression, cells were permeabilized, stained with Alexa 647-conjugated anti-myc antibody, and mean Alexa 647 signal was measured using a BD FACSCalibur flow cytometer.

### cAMP measurements

For all cAMP measurements, a competitive ELISA cAMP assay was used (Enzo Cat #581001 or VWR Cat #75817-364). Cells expressing bPAC constructs were plated on 6-well plates, stimulated with light, and lysed by addition of 0.1 M HCl. In experiments where inhibitors were used to block phosphodiesterase activity, 100 μM IBMX and 10 μM Rolipram were added to the dishes immediately prior to photostimulation. In experiments where Dyngo-4a was used to block endocytosis, 30 μM Dyngo-4a was added to the dishes for 15 min prior to stimulation The ELISA cAMP assay was performed following manufacturer recommendations, and values were normalized to total protein concentration for each respective sample.

### qPCR analysis of target gene expression

bPAC signaling was stimulated with light, then cells were incubated in the dark for 30 min. Total RNA was extracted from the samples using the RNeasy MinElute Cleanup Kit (Qiagen, Cat #74204). Reverse transcription was carried out using Superscript III RT enzyme (Invitrogen, Cat #18080044) following recommended manufacturer protocols. Power SYBR Green PCR MasterMix (ThermoFisher Scientific, Cat #4367659) and the following primers were used for the qPCR reactions- *GAPDH*: F 5’-CAATGACCCCTTCATTGACC-3’ and R 5’-GACAAGCTTCCCGTTCTCAG-3’; *PCK1*: F 5’-CTGCCCAAGATCTTCCATGT-3’ and R 5’-CAGCACCCTGGAGTTCTCTC-3’, and quantified transcript levels were normalized to *GAPDH*.

### SILAC labeling and bPAC stimulation

Cells stably expressing bPAC were grown to <80% confluence in 10-cm round cell culture dishes containing 10 ml of either “Light”-isotope containing medium (lysine- and arginine-depleted medium supplemented with regular lysine and arginine) or “Heavy”-isotope containing medium (supplemented with (13C) lysine and (13C,15N) arginine) for > 6 cell divisions. Two dishes (one “Light”, one “Heavy”) were used for each stimulation replicate. Prior to photostimulation, cells were washed once in PBS and twice in serum-free DMEM, and then grown in 10 ml serum-free DMEM for >16 hrs. Cells grown continuously in the dark served as unstimulated controls. To minimize the impact of SILAC-based labeling artifacts, SILAC medium swap experiments were done for a total of two biological replicates per bPAC cell line, where in replicate #1, “heavy”-medium labeled cells were photostimulated and “light”-medium labeled cells were left in the dark, and in replicate #2, “light”-medium labeled cells were photostimulated and “heavy”-medium labeled cells were left in the dark. Stimulation with a 5-min light pulse was carried out inside a tissue culture incubator, then cells were incubated in the dark for 5 min for a total of 10 min from start of photostimulation until lysis.

### Sample preparation for mass spectrometry

At the end of the 10-min interval, cells were lysed directly in 5 M Urea/0.2% N-dodecyl-maltoside, and phosphatase inhibitors (Sigma phosphatase inhibitor 2 and 3), then sonicated using a Fisher sonicator at 12% amplitude for total of 20 seconds, alternating 10 s on, 10 s off, 10 s on, until lysates were clear. Prior to mixing, approximate concentration was estimated with a Nanodrop, and stimulated vs unstimulated samples were mixed at that point at a 1:1 ratio. Mixed samples were first reduced for 30 minutes with 10 mM TCEP, then alkylated for 30 minutes with 18 mM iodoacetamide and quenched with 18 mM DTT. Prior to trypsin digest, final urea concentration was adjusted to 2 M, then samples were digested overnight at 37°C on a rotator with modified trypsin (1:20 trypsin:sample ratio) (Promega). Peptides were desalted using SepPak C18 columns (Waters) and lyophilized to dryness in a speed-vac. Phosphopeptide enrichment was carried out as previously described using in-house generated Fe^3+^-IMAC resin ^13^. Briefly, 1 mg of dried peptides were resuspended in 80% MeCN/0.2 % TFA and bound to Fe^3+^-IMAC resin. Beads were washed four times in 80% MeCN/0.1% TFA and twice in 0.5% formic acid, and phosphopeptides were eluted in 50% MeCN/0.2% formic acid, dried using speed-vac, and resuspended in 0.1% formic acid for LC/MS analysis.

### Mass spectrometry and data analysis

Purified phosphopeptides resuspended in 0.1% formic acid were analyzed on a Thermo Scientific LTQ Orbitrap Elite mass spectrometry system equipped with a Proxeon Easy nLC 1000 ultra high-pressure liquid chromatography and autosampler system. All samples were analyzed in technical duplicates. Samples were injected onto a C18 column (25 cm x 75 um I.D. packed with ReproSil Pur C18 AQ 1.9 um particles) and subjected to a 4-hour gradient from 0.1% formic acid to 30% ACN/0.1% formic acid. The mass spectrometer collected one full scan at 120,000 resolution in the Orbitrap followed by 20 collision-induced dissociation MS/MS scans for the 20 most intense peaks from the full scan in the dual linear ion trap. Dynamic exclusion was enabled for 30 seconds with a repeat count of 1. Charge state screening was employed to reject analysis of singly charged species or species for which a charge could not be assigned. Andromeda search engine was used within MaxQuant software package (version 1.2.2.5)^31^to align the raw data files against a human protein sequence database downloaded from SwissProt/UniProt (version 03/06/2012). A total of 20,247 entries within the database were searched. Methionine oxidation, protein N-terminus acetylation, and serine, threonine, and tyrosine phosphorylation were set as variable modifications, and cysteine carbamidomethylation was specified as a fixed modification. MaxQuant was configured to generate and search against a reverse sequence database for false discovery rate calculations. The first search was performed with a mass accuracy of +/- 20 parts per million and the main search was performed with a mass accuracy of +/- 6 parts per million. Parameters were set as follows: 1) maximum 5 modifications per peptide, 2) maximum 2 missed cleavages per peptide, 3) maximum peptide charge of 7+. For MS/MS matching, the minimum peptide length was set to 7 amino acids, and the following were allowed: 1) higher charge states, water and ammonia loss events, 2) a mass tolerance of 0.5 Da, and the top 6 peaks per 100 Da were analyzed. Only proteins and peptides falling below a false discovery rate of 1% were considered. Results were matched between runs with a time window of 2 minutes for technical duplicates **(Dataset S3)**. The data were condensed by a custom Perl script that takes the maximum intensity of any unique peptide and charge state between the two technical replicates, log-transformed using log base 2, and median-centered. For the “heavy”-labeled unstimulated/”light”-labeled photostimulated samples, the inverse values for all original log_2_ ratios were taken to aid the ease of averaging the biological replicates. To identify “cAMP target phosphopeptides”, we considered peptides with log_2_ values with 2 standard deviations above or below the sample mean in each of two Cyto-bPAC replicates **(Dataset S1)**. Protein localization from the Human Protein Atlas^32^, inferred from antibody-based immunofluorescence microscopy, was manually curated further based on published reports. Significantly enriched Gene Ontology categories were identified with Panther^33^ **(Dataset S1)**, and protein-protein interaction networks were generated using String database^34^ based on biochemical data and annotated interactions from curated databases with a threshold confidence score of 0.400. The resulting network data were visualized based on averaged Cyto-bPAC log_2_ values using the Cytoscape software^35^. Enrichment of amino acid motifs was determined with MotifX software^36^in R using statistical cut-off *p* < 1.0×10^−6^ by Fisher’s exact test and minimal number of occurrences= 20. Average linkage hierarchical clustering was performed with the Cluster software^37^ using Euclidian distance as a similarity metric and visualized with Java TreeView^38^. To elucidate differentially modified phosphopeptides between sets of conditions, we considered only cAMP target phosphopeptides measured in all biological replicates for each pairwise comparison, and analyzed by multiple t-test using the adaptive Benjamini-Hochberg step-up procedure^39^, and FDR of 10% was used as cut-off **(Dataset S2)**. To examine if there are location-specific differences in the distribution of upregulated and downregulated phosphosites, we carried out linear regression analysis of the abundance values between pairs of conditions, computed the residuals for the regression, and used unpaired t-test analysis assuming that the populations have the same standard deviation.

### Western blotting and quantification

Cells were grown in 6-well dishes, and medium was switched to serum-free overnight. GPCR internalization was acutely inhibited by treatment of cells with 30 μM Dyngo-4A for 15 min. Cells were then stimulated with 1 μM isoproterenol or 100 nM AVP for 10 min, lysed in ice-cold Lysis Buffer (50 mM Tris, 150 mM NaCl, 1% Triton X-100, 0.5% Sodium Deoxycholate, 0.1% Sodium Dodecyl Sulfate, protease inhibitors cocktails). Lysates were quantified and equal amounts of total protein were loaded on a protein gel. Nitrocellulose membranes were visualized using the Odyssey imager system (Li-COR) and individual bands (Figure S3b,d) were quantified. Phospho-PKA bands were normalized to the loading control GAPDH.

## Data availability

Mass spectrometry data are available via ProteomeXchange with identifier PXD025775.

## Acknowledgements

We thank members of the von Zastrow and Krogan laboratories for discussions and critical feedback on the manuscript. We also thank Angus Nairn (Yale University, CT) for advice on protein phosphatase 2, and the Lefkowitz laboratory (Duke University, NC) for generously providing the V2R construct.

## Author contributions

N.G.T and M.v.Z. designed the study and interpreted the results. N.G.T., M.T-Z. and B.W.N. performed the phosphoproteomic experiments. N.G.T. analyzed the data. G.E.P. performed the Western blot experiments. N.J.K. provided access to mass spectrometry equipment and reagents. J.R.J., D.J-M., A.P.K. and B.W.N. contributed mass spectrometry analysis tools and carried out data upload. N.G.T. and M.v.Z. wrote the manuscript. All authors edited the manuscript.

## Funding and additional information

This work was supported by the National Institute of Health (Grants MH109633 to N.G.T; HL129689 to G.E.P.; DA010711 and DA012864 to M.v.Z.; GM081879, GM082250, GM107671, and U19 AI106754 to N.J.K.). G.E.P. was also supported by the American Heart Association (16PRE26420057). The content is solely the responsibility of the authors and does not necessarily represent the official views of the National Institutes of Health.

## Conflict of interest

The authors declare that they have no conflicts of interest with the contents of this article.

## List of abbreviations/keywords

cAMP: cyclic AMP
bPAC: bacterial photoactivatable adenylyl cyclase
GPCR: G protein-coupled receptor
SILAC: stable isotope labeling with amino acids in cell culture
Fe^3+^: NTA IMAC-iron (III)-nitrilotriacetic acid immobilized metal ion affinity
LC: MS/MS-liquid chromatography-tandem mass spectrometry
PKA: protein kinase A
PDE: phosphodiesterase
PP2A: protein phosphatase 2A

---

**Figure S1.**
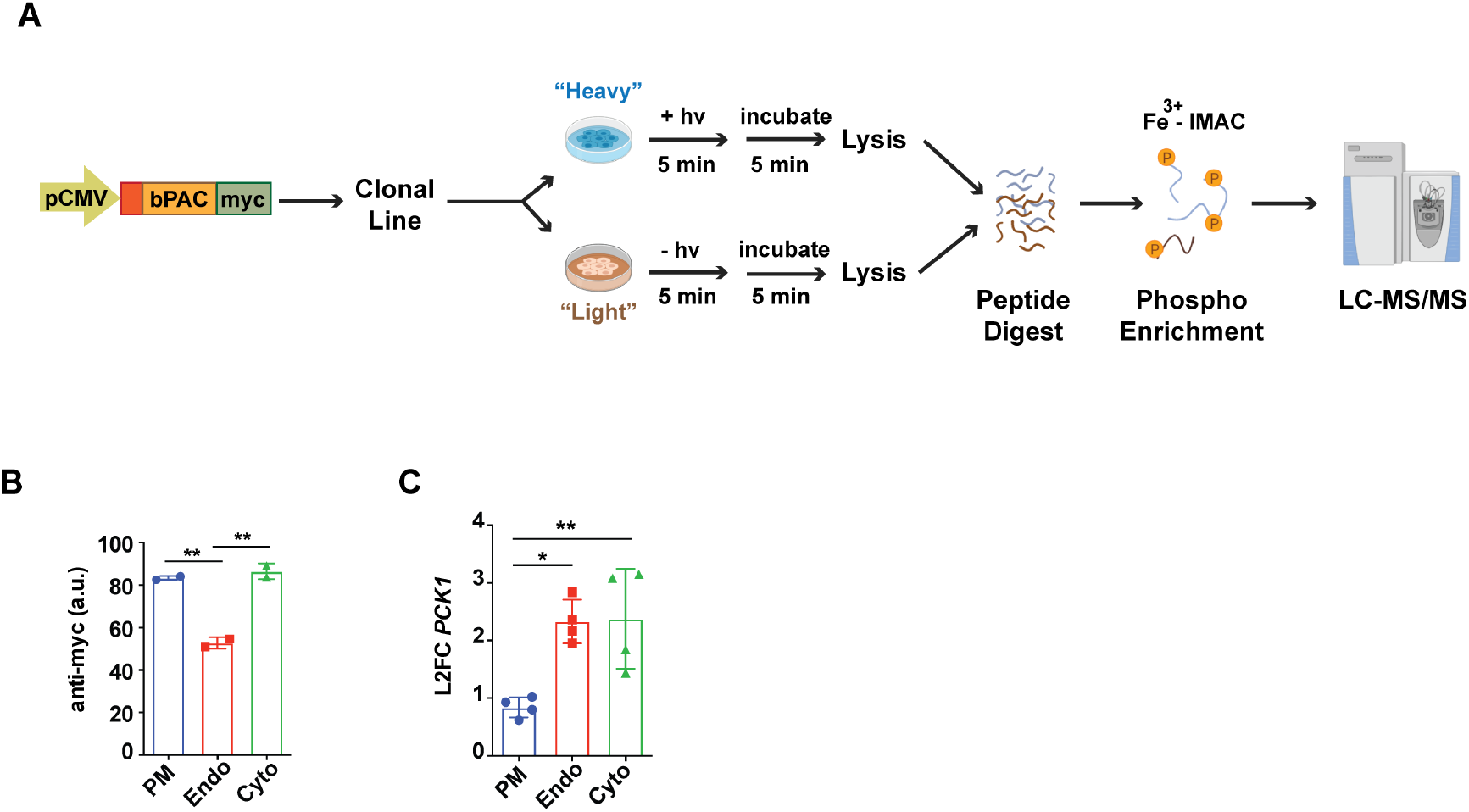
Characterization of bPAC expression and downstream signaling responses in the stable cell lines. **(a)** Schematic overview of the experimental workflow. Red square denotes an organelle-targeting motif (Lyn or 2xFYVE). Created with BioRender.com. **(b)** bPAC expression levels quantified by flow cytometry analysis with Alexa 647-conjugated anti-myc antibody in fixed cells. Data are mean of n = 2. **(c)** RT-qPCR quantification of *PCK1* transcriptional upregulation upon bPAC photostimulation. Expression levels were normalized to a housekeeping gene (*GAPDH*), and displayed as fold-change relative to unstimulated cells. Data are mean of n = 4. Error bars = ± SD. ** = *p* ≤ 0.01, * = *p* ≤ 0.05 by one-way ANOVA test, using Tukey’s multiple testing correction. PM = plasma membrane, Endo = early endosome, Cyto = cytosol.

**Figure S2.**
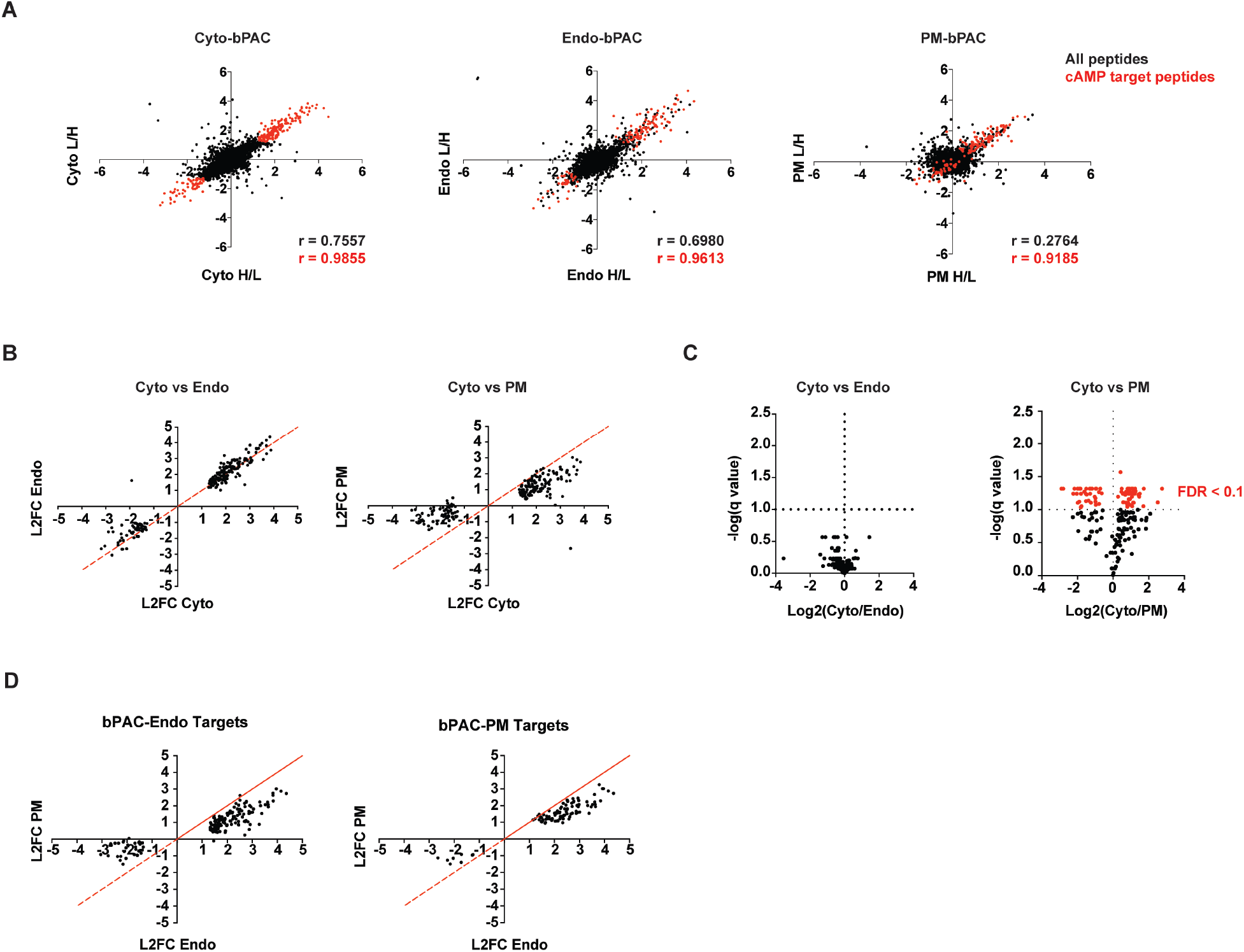
SILAC-mass spectrometry analysis of global phosphoresponses to cAMP production. **(a)** Scatter plots depicting phosphopeptide abundance correlations between biological replicates. For each experimental condition, a pair of SILAC isotope label-swapped photostimulated vs unstimulated cells was included. Log-transformed median-normalized fold changes are plotted on the axes. For the L/H (“light” vs “heavy” isotope-labeled) replicates corresponding to unstimulated/stimulated log_2_ ratios (L2FC), the inverse values for the original ratios are shown for convenience. Black = all quantified phosphopeptides, red = cAMP target phosphopeptides defined based on the bPAC-Cyto experiments **(Dataset S1)**; corresponding Pearson correlation coefficients are color-coded. **(b)** Scatter plots comparing the abundance of cAMP target phosphopeptides in bPAC-Endo, bPAC-Cyto, and bPAC-PM cells. Red line denotes x = y. **(c)** Volcano plots showing pairwise comparisons between the changes in cAMP-dependent phosphopeptide abundance relative to respective unstimulated conditions. In red = significantly different phosphopeptides summarized in **Dataset S2** (FDR ≤ 10%). **(d)** Scatter plots comparing the abundance of target phosphopeptides defined based on bPAC-Endo (left) or bPAC-PM (right) experiments. Sites were defined as targets if their corresponding phosphopeptides displayed abundance changes in response to photostimulation with z-scores ≥ 2 or ≤ −2 in each Endo or PM replicate, respectively. Red line denotes x = y. Data in this Figure are average from n = 2.

**Figure S3.**
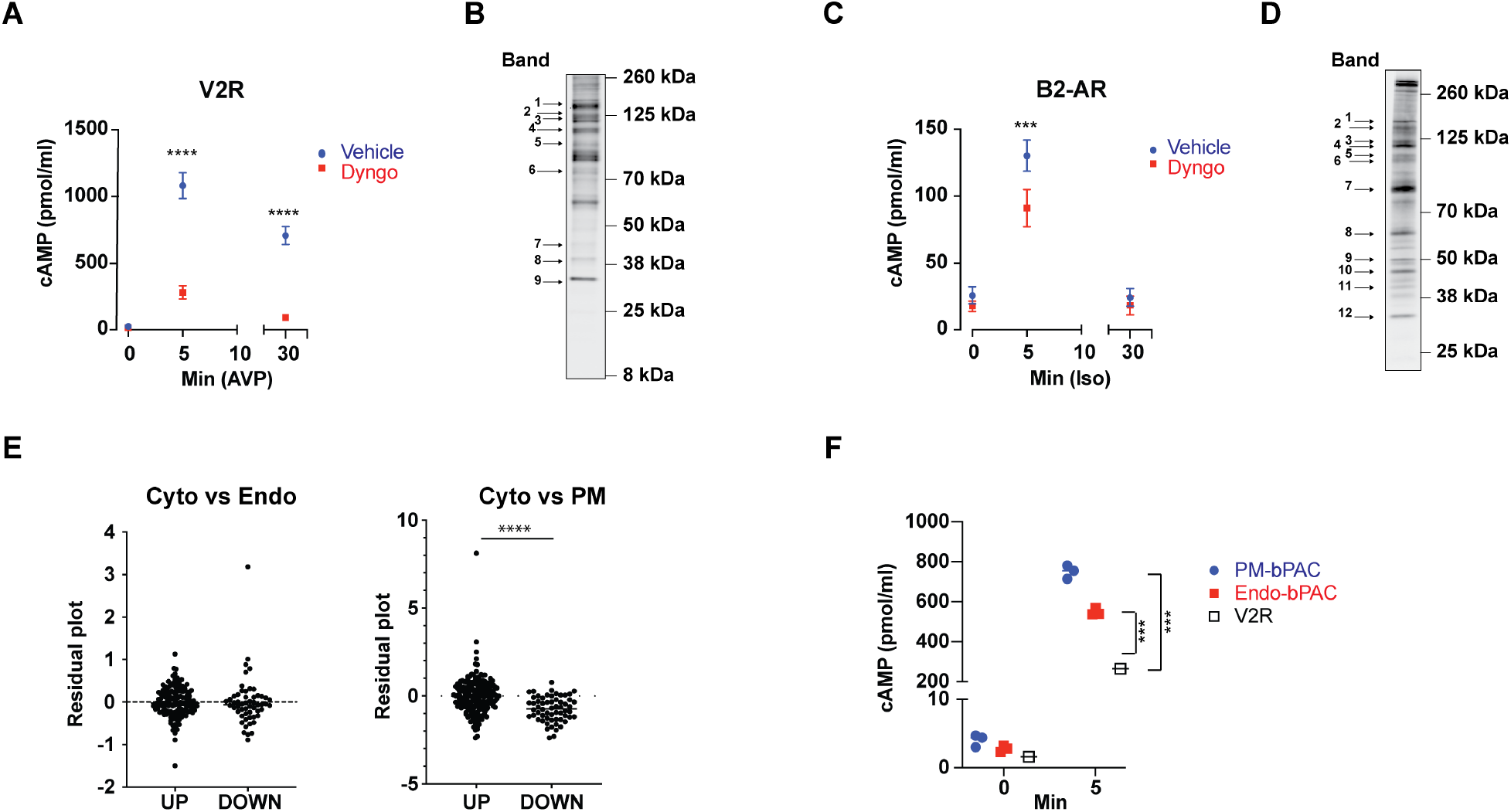
Impact of cAMP localization on the cellular phosphoresponses. **(a)** Endocytic blockade diminishes cAMP production downstream of the vasopressin receptor (V2R). Cells were pre-treated with 30 μM Dyngo-4a or vehicle (DMSO) for 15 min, then stimulated with 100 nM arginine vasopressin (AVP) for indicated times. Total cAMP was measured by ELISA assay, and all values were normalized to total protein per sample. Data are mean from n = 3. **(b)** Western blot bands included in quantification analysis in **Fig. 2d**. **(c)** Endocytic blockade diminishes cAMP production downstream of the beta2-adrenergic receptor (β2-AR). Cells were pre-treated with 30 μM Dyngo-4a or vehicle (DMSO) for 15 min, then stimulated with 1 μM isoproterenol (Iso) for indicated times. Total cAMP was measured by ELISA assay, and all values were normalized to total protein per sample. Data are mean from n = 3. **(d)** Western blot bands included in quantification analysis in **Fig. 2e**. **(e)** Residual plots computed from linear regression analyses on datasets from bPAC-Endo, bPAC-Cyto and bPAC-PM cells. Residuals are shown for peptides with upregulated versus downregulated phosphorylations. Data are mean from n = 2. **(f)** Total cAMP produced following stimulation of the bPACs with light or the V2R with 100 nM AVP, measured by ELISA cAMP assay and normalized to total protein per sample. Data are mean from n = 3. Error bars = ± SD. **** = *p* ≤ 0.0001, *** = *p* ≤ 0.001, ** = *p* ≤ 0.01 by two-way ANOVA test using Sidak’s correction in **(a,c,f)** or unpaired two-sided t-test in **(e)**.

## Notes

### Competing Interest Statement

The authors have declared no competing interest.

